# Bacterial protein interaction networks: connectivity is ruled by gene conservation, essentiality and function

**DOI:** 10.1101/681395

**Authors:** Maddalena Dilucca, Giulio Cimini, Andrea Giansanti

**Affiliations:** Dipartimento di Fisica, Sapienza University of Rome, Rome, Italy; IMT School for Advanced Studies, Lucca, Italy; Istituto dei Sistemi Complessi (ISC)-CNR, Rome, Italy; INFN Roma1 unit, Rome, Italy

## Abstract

Protein-protein interaction (PPI) networks are the backbone of all processes in living cells. In this work we study how conservation, essentiality and functional repertoire of a gene relate to the connectivity *k* of the corresponding protein in the PPI networks. Focusing on a set of 42 mostly distantly related bacterial species, we investigate three issues: i) whether the distribution of connectivity values changes between PPI subnetworks of essential and nonessential genes; ii) how gene conservation, measured both by the evolutionary retention index (ERI) and by evolutionary pressure (as represented by the ratio *K*_*a*_*/K*_*s*_) is related to the the connectivity of the corresponding protein; iii) how PPI connectivities are modulated by evolutionary and functionaly relationships, as represented by the Clusters of Orthologous Proteins (COGs). We show that conservation, essentiality and functional specialization of genes control in a universal way the topology of the emerging bacterial PPI networks. Noteworthy, a structural transition in the network is observed such that, for connectivities *k* ≥ 40, bacterial PPI networks are mostly populated by genes that are conserved, essential and which, in most cases, belong to the COG cluster J, related to ribosomal functions and to the processing of genetic information.

## Introduction

To operate biological activities in living cells, proteins work in association with other proteins, possibly assembled in large complexes. Hence, knowing the interactions of a protein is important to understand its cellular functions. Moreover, a comprehensive description of the stable and transient protein-protein interactions (PPIs) within a cell would facilitate the functional annotation of all gene products, and provide insight into the higher-order organization of the proteome [1, 2]. Several methodologies have been developed to detect PPIs, and have been adapted to chart interactions at a proteome-wide scale. These methods, that combine different technologies with complementary experiments and computational analyses, were shown to generate high-confidence PPI networks, enabling the assignment of several proteins to functional categories [3, 4].

Moreover, the statistical study of bacterial PPIs over several species (meta-interactomes) has brought important knowledge about protein functions and cellular processes [5, 6]. This work contributes to this line of research. Our aim is to shed light on the relationship between conservation, essentiality and functional annotation at the genetic level with the connectivity patterns of the PPI networks. We extend previous observations which suggested a strong correlation between codon bias and the topology of PPI networks on the one hand, and between codon bias and gene conservation/essentiality on the other hand [7, 8]. It is worth, in the next two paragraphs, to make more precise what is usually meant bygene essentiality and gene conservation.

Individual genes in the genome contribute differentially to the survival of an organism. According to their known functional profiles and based on experimental evidence, genes can be divided into two categories: essential and nonessential ones [9, 10]. Essential genes are not dispensable for the survival of an organism in the environment it lives in [10, 11]. Nonessential genes are instead those which are dispensable [12], being related to functions that can be silenced without compromising the survival of the organism. Naturally, each species has adapted to one or more evolving environments and, plausibly, genes that are essential for one species may be not essential for another one.

It has been argued many times that essential genes are more conserved than nonessential ones [13–17]. The term “conservation” has however at least two meanings. On the one hand, a gene is conserved if orthologous copies are found in the genomes of many species, as measured by the Evolutionary Retention Index (ERI) [9, 18]. On the other hand, a gene is (evolutionarily) conserved when it is subject to a purifying evolutionary pressure which disfavors mutations. This pressure is usually measured by *K*_*a*_*/K*_*s*_, the ratio of the number of nonsynonymous substitutions per nonsynonymous site to the number of synonymous substitutions per synonymous site. In this second meaning a conserved gene is, in a nutshell, a slowly evolving gene [13, 19].

In this work we show that bacterial PPI networks display an interesting topological-functional transition, ruled by protein connectivity *k* and with a threshold between *k* = 40 and *k* = 50. Proteins with high PPI network connectivities (hubs) likely correspond to genes that are conserved and essential. Conversely, genes that correspond to hub proteins in the PPI network are likely to be essential and conserved. Additionally, below the threshold the functional repertoire of proteins is heterogeneous, whereas, above the threshold there is a quite strict functional specialization.

## Materials and Methods

We consider a set of 42 selected bacterial genomes (that we have previously investigated in [8]), reported in Table 1. These species were chosen in order to have the largest possible coverage of available data for what concerns conservation, essentiality and selective pressure of genes. Nucleotide sequences were downloaded from the FTP server of the National Center for Biotechnology Information. [20].

**Table 1.**
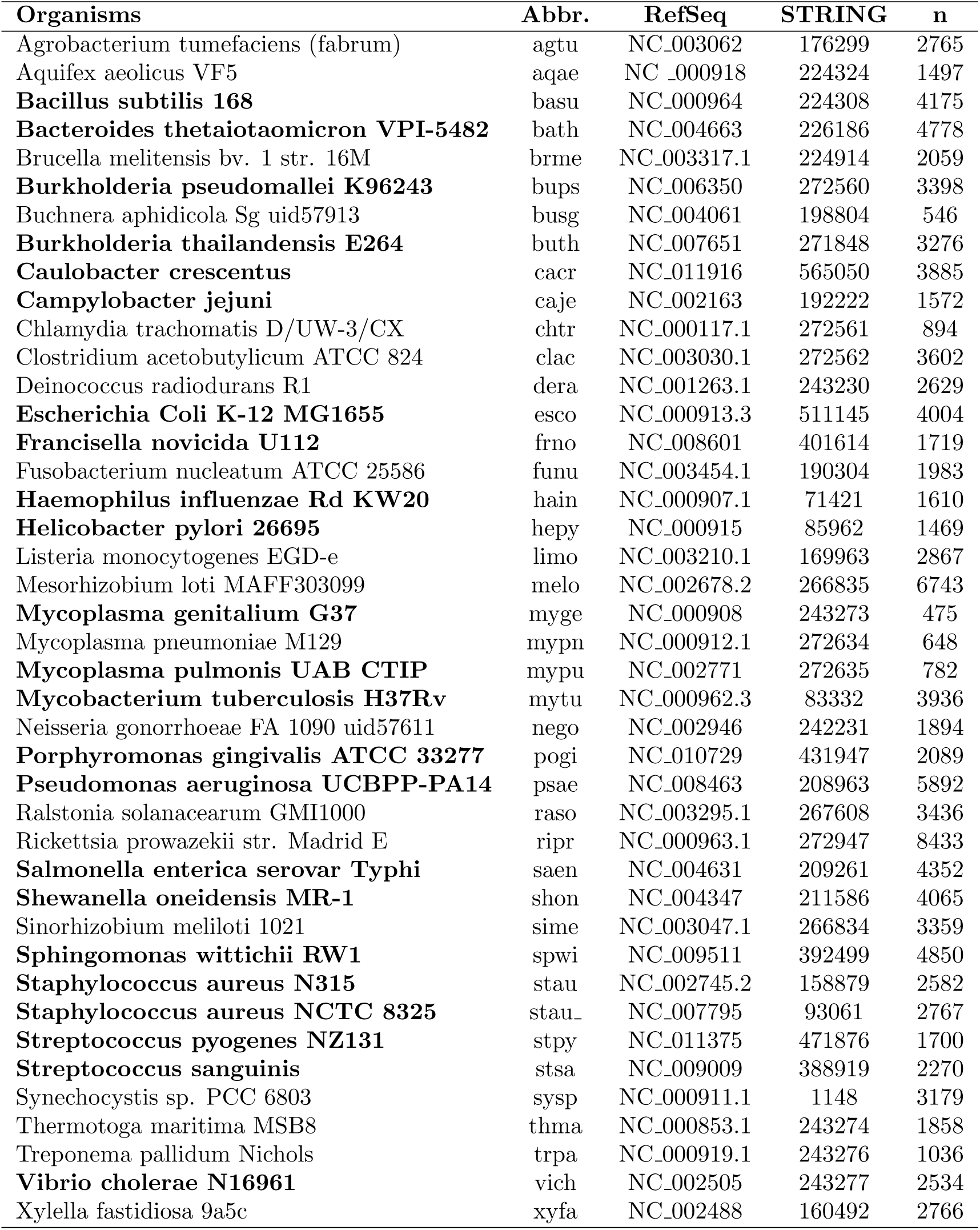
Summary of the selected bacterial dataset. Organism name, abbreviation, RefSeq, STRING code, size of genome (number of genes *n*). Genomes annotated in the Database of Essential Genes (DEG) are highlighted with bold fonts.

### Gene Conservation

We use the Evolutionary Retention Index (ERI) [9] as a first way of measuring conservation of a gene. The ERI of a gene is the fraction of genomes (among those reported in Table 1) that have at least an ortholog of the given gene. Then, a low ERI value means that a gene is specific, common to a small number of genomes, whereas high ERI is a characteristic of highly shared, putatively universal and essential genes.

We also make reference to another notion of gene conservation. Conserved genes are those which are subject to a purifying, conservative evolutionary pressure. To discriminate between genes subject to purifying selection and genes subject to positive selective Darwinian evolution, we use a classic but still widely used indicator, the ratio *K*_*a*_/*K*_*s*_ between the number of nonsynonymous substitutions per nonsynonymous site (*K*_*a*_) and the number of synonymous substitutions per synonymous site (*K*_*s*_) [19]. Conserved genes are characterized by *K*_*a*_*/K*_*s*_ < 1. We used *K*_*a*_*/K*_*s*_ estimates by Luo [15] that are based on the method by Nej and Gojobori [21].

### Gene Essentiality

We used the Database of Essential Genes (DEG, *www.essentialgene.org*) [15], which classifies a gene as either essential or nonessential on the basis of a combination of experimental evidence (null mutations or trasposons) and general functional considerations. DEG collects genomes from Bacteria, Archea and Eukarya, with different degrees of coverage [22, 23]. Of the 42 bacterial genomes we consider, only 23 are covered—in toto or partially—by DEG, as indicated in Table 1.

### Protein-Protein Interaction Network

PPIs are obtained from the STRING database (Known and Predicted Protein-Protein Interactions, *https://string - db.org/*) [24]. We have chosen STRING because it is the only available database with 1678 bacteria species, thus useful to extend analysis such as those performed in [7] to multiple species. The problem with STRING (functional associations might be used to represent both physical protein-protein molecular interactions and more abstract meta-interactome interactions with it our best to modulate our results using the parameter *w*, offered by STRING to its users. In STRING, each interaction is assigned with a confidence level or probability *w*, evaluated by comparing predictions obtained by different techniques [25–27] with a set of reference associations, namely the functional groups of KEGG (Kyoto Encyclopedia of Genes and Genomes) [28]. In this way, interactions with high *w* are likely to be true positives, whereas, a low *w* possibly corresponds to a false positive. As usually done in the literature, we consider only interactions with *w* ≥ Θ and select a stringent cut-off Θ = 0.9 that allows for a fair balance between coverage and interaction reliability (see for instance the case of *E.coli* reported in [7]).

We denote by *k* the *degree* (number of connections) associated to each proteins in each PPI network after the thresholding procedure. Note also that after applying the cut-off we are left, for each network, with a number of isolated proteins (with no connections) that grows as 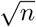 (where *n* is the number of proteins in the genome). These proteins are not considered in the network analysis and are regarded as stemming from statistical noise.

### Clusters of orthologous proteins

We use functional annotation given by the popular database of orthologous groups of proteins (COGs) from Koonin’s group, available at *http://ncbi.nlm.nih.gov/COG/* [29, 30]. We consider 15 functional COG categories (see Table 2), excluding the generic categories R and S for which functional annotation is too general or missing.

**Table 2.**
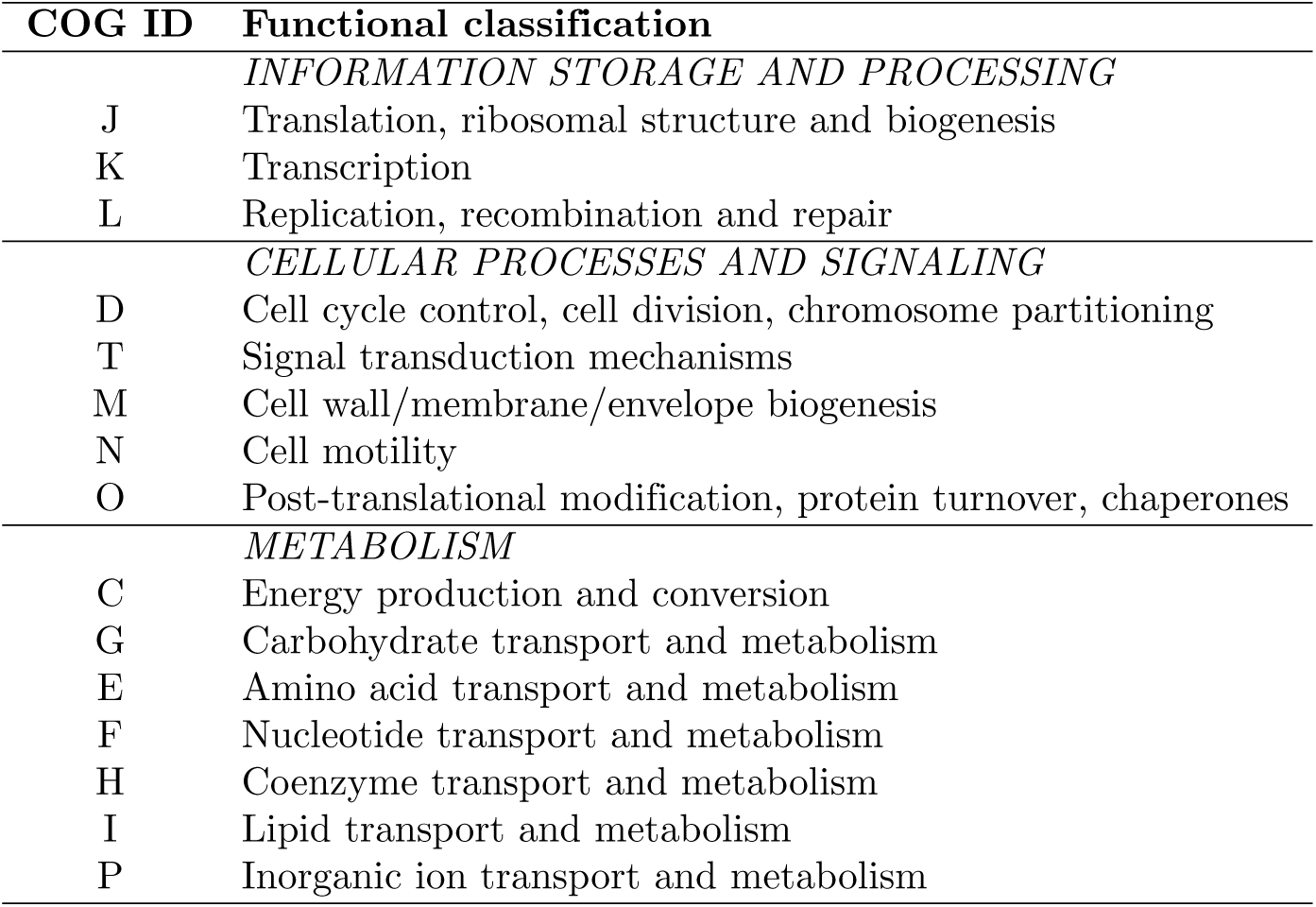
Functional classification of COG clusters.

## Results and discussion

### Degree distribution of PPI networks

We start by studying the degree distributions *P* (*k*) observed in bacterial PPIs. We first recall that such a distribution was found to be scale-free in *E.coli* [7, 31–33], meaning that the corresponding PPI network features a large number of poorly connected proteins, and a relatively small number of highly connected hubs. In order to assess the generality of this observation, we compute *P* (*k*) for each genome of Table 1 (plots are reported in figures S2-S3 of the Supplementary Information). Note that, despite the fact that PPIs of different bacteria have different sizes and densities, their average connectivity and the support of their *P* (*k*) are very similar (as shown in figure S1 of the Supplementary Information). Thus, we can superpose all the considered bacterial degree distributions without the need to normalise the support of each *P* (*k*). When doing so, we observe two distinct regimes (see figure 1). For low values of *k*, the distribution has a scale-free shape *P* (*k*) *∝ k*^−*γ*^. This result is consistent with previous findings for yeast, worm and fly [34]and for co-conserved PPIs in some bacteria [35]. For higher values of *k*, however, the distribution deviates from a power law, and a bump with a Gaussian-like shape emerges. Interestingly, this feature is almost undetectable taking individual species alone as *E.coli* (see fig 2 in supporting materials of Dilucca [7]), but clearly emerges when the statistics is enriched by adding together several species. The bump emerging for *k* ≥ 40 is reasonably due to the contribution of proteins belonging to complexes [36]. Indeed, if we consider the separate contribution of essential and nonessential genes to the *P* (*k*) (for DEG-annotated genomes), we see that the superposed peak is present only in the degree distribution of essential genes. Moreover, the degree distributions for essential and nonessential genes are well separated and the average degree is systematically higher for essential genes than for nonessential ones—consistently with previous findings [34].

**Fig 1.**
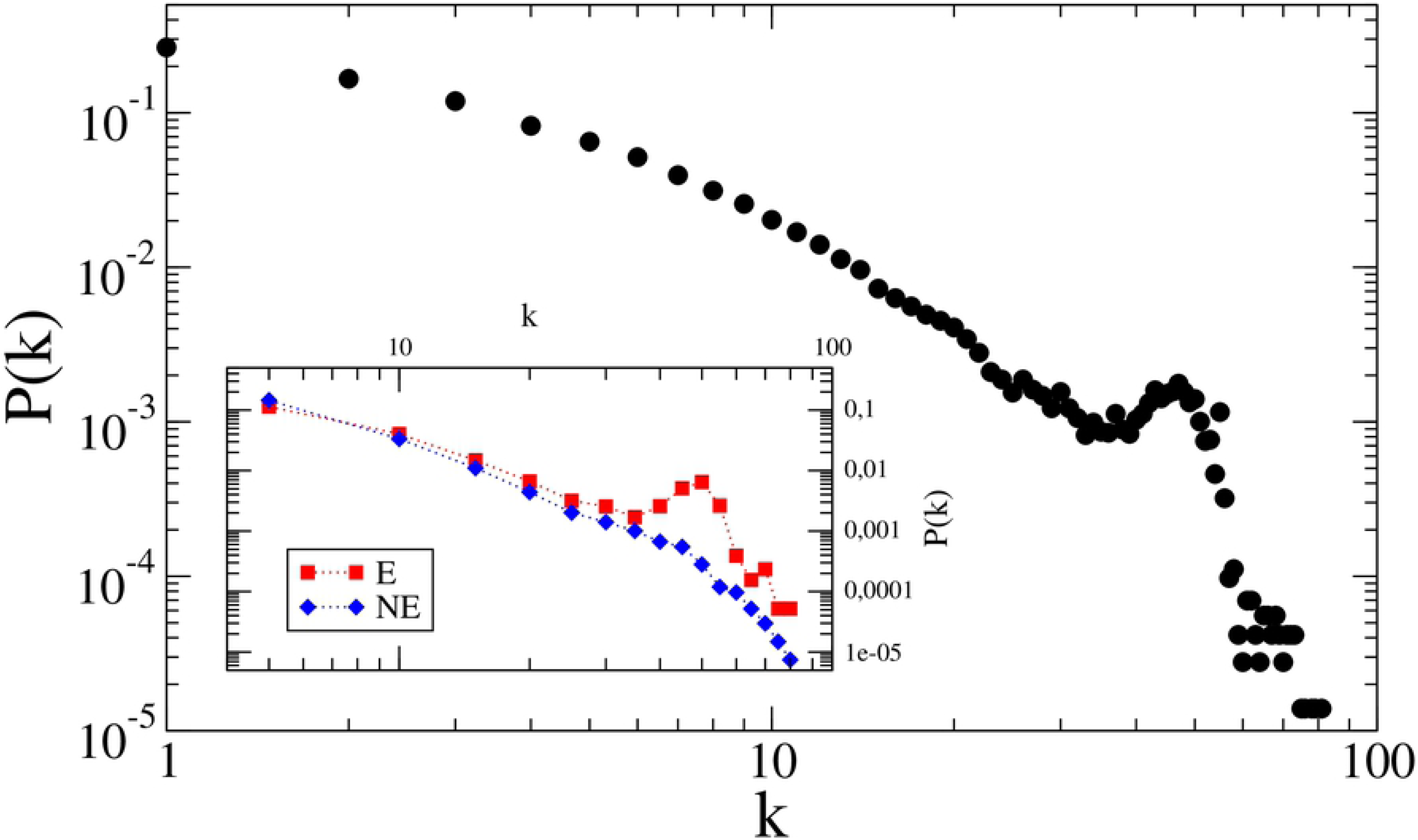
Probability distribution *P* (*k*) for the number of connections *k* of each protein, averaged over the bacterial species considered in Table 1. Inset: *P* (*k*) for essential (E) and nonessential (NE) genes, averaged over DEG-annotated genomes. Note that the average degree is higher for essential genes than for nonessential ones, and the two probability distributions are quite distinct. The region of the curve for low *k* can be well approximated by a power law [37].

**Fig 2.**
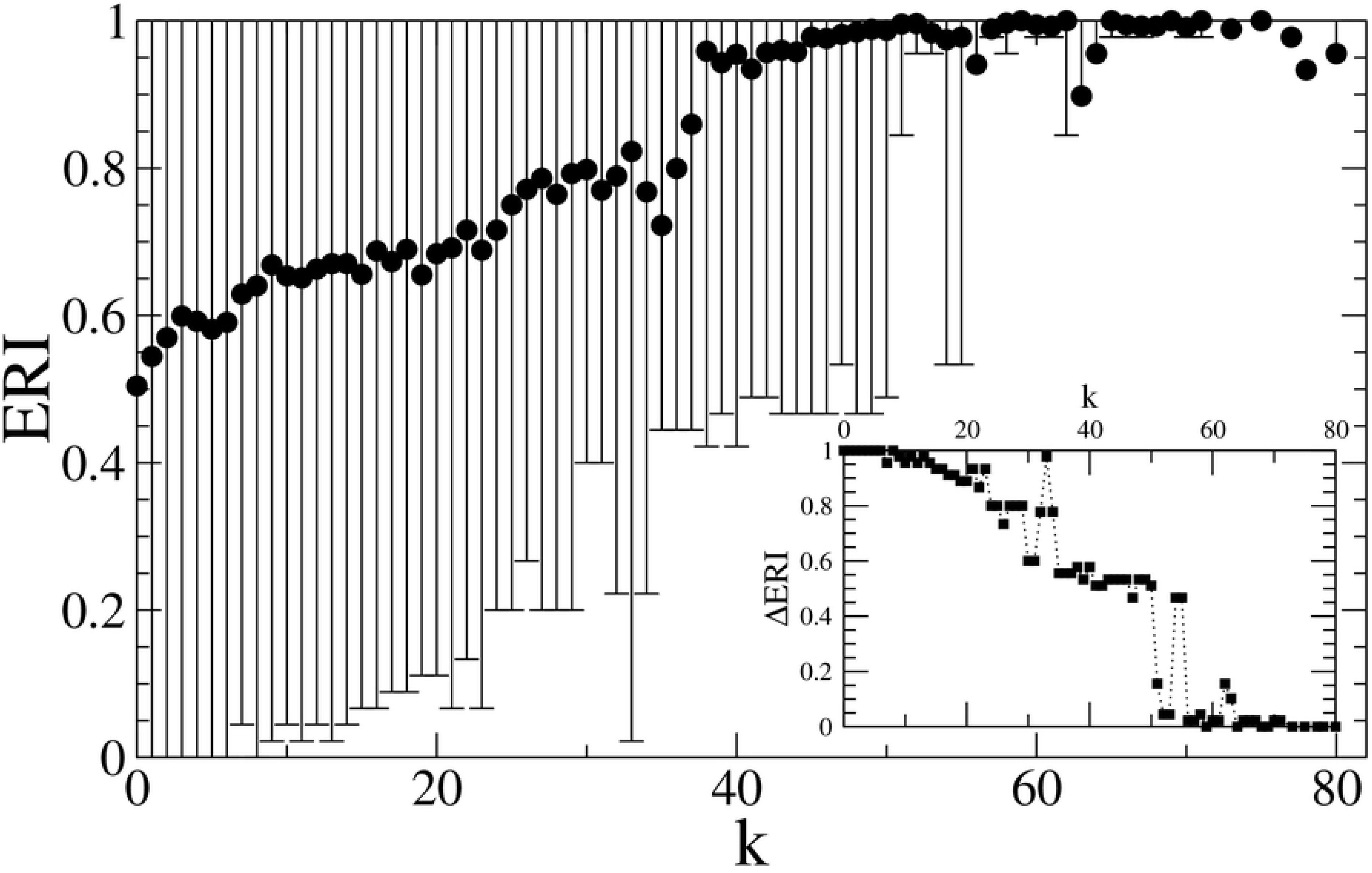
Average ERI values of bacterial genes as a function of the degrees *k* of the corresponding proteins, for all the considered genomes. Error bars are standard deviations of ERI values associated to a given *k* value. Inset: amplitude of the error bar (ΔERI) as a function of *k*.

### PPI connectivity and gene conservation

We now investigate whether there is a correlation between the degree of conservation (as measured by ERI) of genes and the connectivity *k* of the corresponding proteins in PPI networks. Figure 2 shows that highly connected proteins are also highly conserved among the bacterial species we consider, that constitute a reasonably wide sample of different evolutionary adaptations. This observation is a strong signature of the existence of an invariant structure of conserved hubs in all bacterial PPI networks. Indeed, we observe the existence of a structural transition for bacterial proteins with *k* > 40, which have ERI close to 1 almost surely, and are thus highly conserved among the species. Proteins that are less connected, on the contrary, have a wide range of ERI values. Interestingly, as shown in the inset of figure 2, the fluctuations in ERI as a function of *k* abruptly decrease for connectivities above the threshold.

We then look at the evolutionary pressure exerted on genes whose proteins have different connectivities. The graph in figure 3 shows the ratio *K*_*a*_*/K*_*s*_ for groups of genes binned by the connectivity *k* of the corresponding proteins, for all the 42 bacterial species considered here. We see that the more connected proteins correspond to genes which are subject to an increasing purifying evolutionary pressure. Indeed, values of the *K*_*a*_*/K*_*s*_ systematically decrease until they become zero, as a function of *k*. This result point to the fact that the more proteins are connected in the PPI networks the more the genes encoding them are subject to a purifying evolutionary pressure.

**Fig 3.**
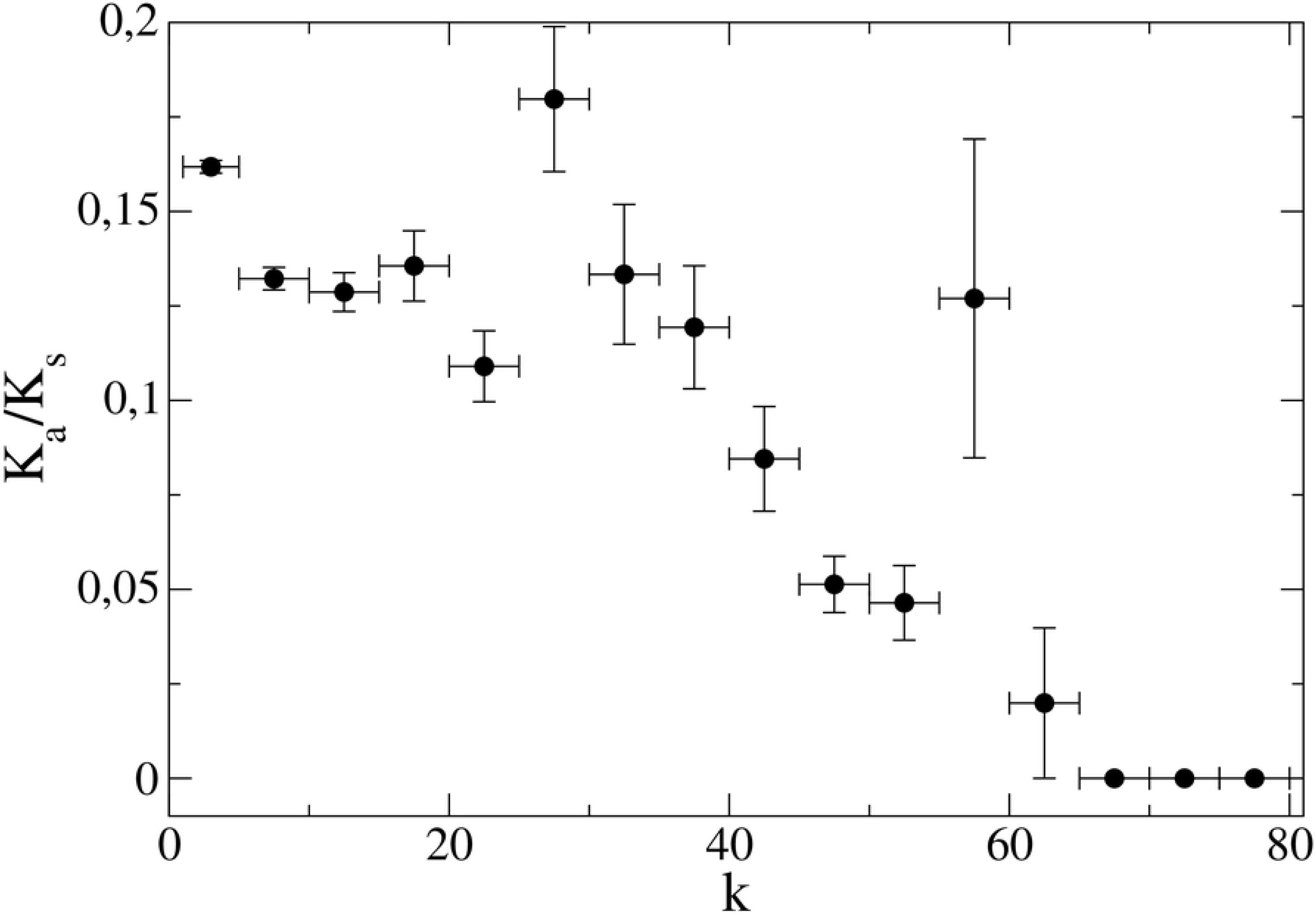
Dependence of selective pressure in terms of *K*_*a*_*/K*_*s*_ for a gene on the degree *k* of the corresponding protein, for all the considered genomes.

### PPI and Essentiality

To further investigate the relationship between gene essentiality and protein connectivities, we consider DEG-annotated genomes and classify interactions between proteins (links) making references to the essentiality of the corresponding genes. We distinguish three sets of links: *ee* (linking proteins from two essential genes), *ēē* (from two nonessential genes) and *eē* (from an essential gene and a nonessential one). We then compute the *density* of these sets of links respectively as:

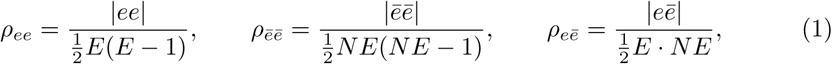

where *E* and *NE* denote the number of essential and nonessential genes, respectively (self-connections are excluded in our analysis). Such densities are then compared with the overall density of the network—restricted to genes classified as either essential or nonessential:

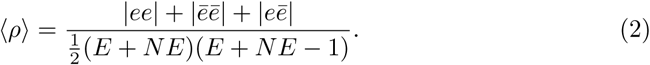

We use the ratios *r*_*ee*_= *ρ*_*ee*_/⟨*ρ*⟩,*r*_*ēē*_= *ρ* _*ēē*_/ ⟨*ρ*⟩ and *r*_*eē*_= *ρ*_*eē*_ /⟨*ρ*⟩ to assess the level of connectivity of the subnetworks with respect to the overall connectivity. Table 3 shows that subnetworks of essential genes are far denser than the overall networks, and that, in general, essential and nonessential genes tend to form network components that are weakly interconnected. This happens because many essential genes encode for ribosomal proteins, which in turn are localized in the ribosome so that they have a major probability to interact [38]. Figures S4-S5 of the Supplementary Information display such network features for each individual species.

**Table 3.** Relative density values r for PPI subnetworks between essential genes (r_*ee*_), between nonessential genes (r_*ēē*_) and between essential and nonessential genes (r_*eē*_), for each DEG-annotated bacterial genome.

| Organisms | $r_{ee}$ | $r_{\bar{e}\bar{e}}$ | $r_{e\bar{e}}$ |
| --- | --- | --- | --- |
| basu | 44.46 | 0.80 | 0.11 |
| bath | 20.07 | 0.76 | 0.25 |
| bups | 6.21 | 0.83 | 0.27 |
| buth | 18.69 | 0.70 | 0.22 |
| cacr | 18.40 | 0.70 | 0.15 |
| caje | 3.65 | 0.82 | 0.32 |
| esco | 2.91 | 0.88 | 0.31 |
| frno | 9.84 | 0.52 | 0.18 |
| hain | 1.65 | 1.15 | 0.27 |
| hepy | 2.91 | 0.78 | 0.38 |
| myge | 1.42 | 0.29 | 0.08 |
| mypu | 3.42 | 0.22 | 0.12 |
| mytu | 8.09 | 0.78 | 0.23 |
| pogi | 11.03 | 0.41 | 0.21 |
| psae | 9.85 | 0.92 | 0.16 |
| saen | 28.80 | 0.81 | 0.12 |
| shon | 6.50 | 0.64 | 0.16 |
| spwi | 15.47 | 0.74 | 0.22 |
| stau | 23.05 | 0.58 | 0.23 |
| stau_ | 21.89 | 0.64 | 0.16 |
| stpy | 9.30 | 0.73 | 0.23 |
| stsa | 30.65 | 0.61 | 0.22 |
| vich | 8.37 | 0.81 | 0.19 |

### PPI connectivity and functional specialization

For each PPI network, we define the conditional probability that a protein with degree *k* belongs to a given COG as:

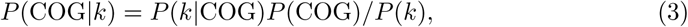

where *P* (*k*) is the degree distribution in the PPI network, *P* (COG) is the frequency of that COG in the proteome, and *P* (*k* | COG) is the degree distribution restricted to that COGs. Figure 4 shows the COG spectrum as a function of *k* over all bacteria species considered. Interestingly, we again note a marked transition. Below *k≃* 40 the COG spectrum is quite heterogeneous: genes corresponding to proteins with low connectivity are spread over several COGs which correspond to different functions (see Table 2). Genes whose proteins are highly connected (*k*≥ 40) are instead mainly concentrated in COG J, which encompasses translation processes and ribosomal functions. There are yet a handful of outliers, hubs with connectivity between 57 and 62, that belong to COG I (related to lipid transport and metabolism) and K and L (which, together with J, define the functional class of information storage and processing). The list of these outliers is reported in Table 4).

**Table 4.**
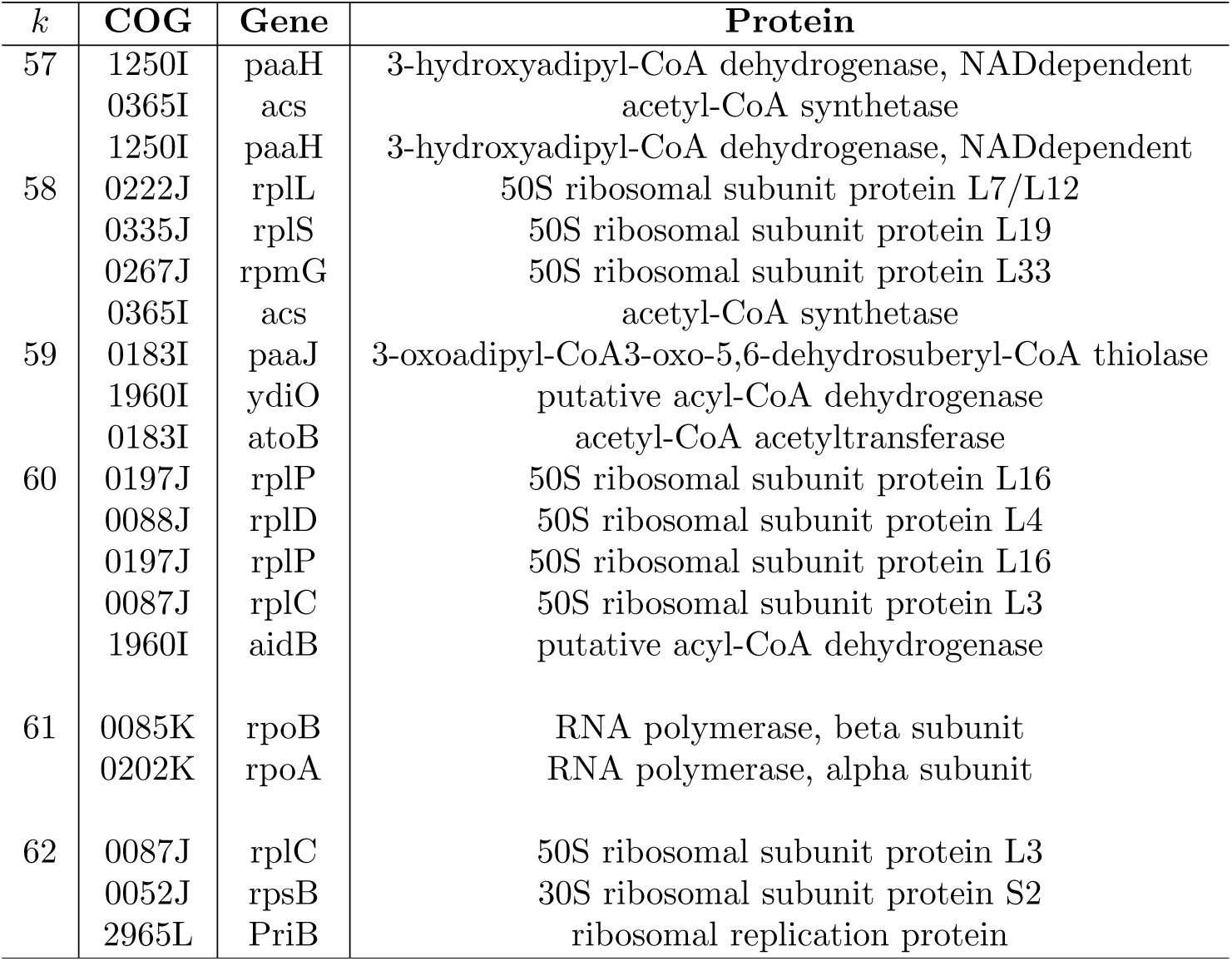
Specific hubs. In this table we detail which proteins populate the few bins of connectivity around *k* = 60 in figure 4.

**Fig 4.**
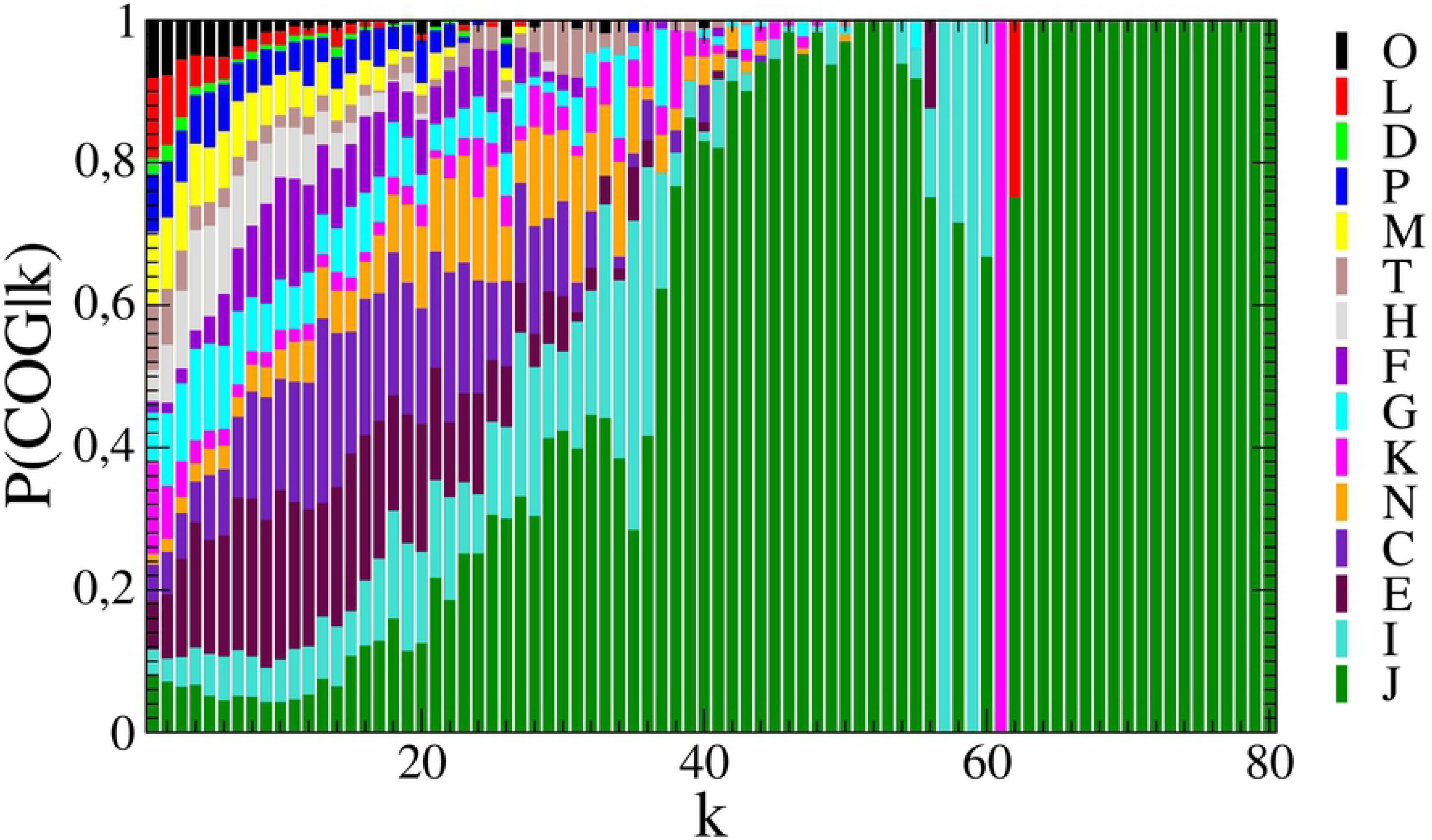
Probability distribution *P* (COG | *k*) of belonging to a given COG for proteins with degree *k*, over all considered genomes. Proteins with low connectivity have a very heterogeneous COG composition, whereas, those with high *k* basically belong only to COG J.

## Conclusions

Topological analysis of biological networks, such as protein-protein interaction or metabolic networks, has demonstrated that structural features of network subgraphs are correlated with biological functions [39, 40]. For instance, it was shown that highly connected patterns of proteins in a PPI are fundamental to cell viability [41]. In this work we have shown the existence of a topological-functional transition in bacterial species, ruled by the connectivity of proteins in the PPI networks. The threshold in *k* of the transition is located between *k* = 40 and *k* = 50. Proteins that have connectivities above the threshold are mostly encoded by genes that are conserved (as measured both by ERI and *K*_*a*_*/K*_*s*_) and essential. Moreover the functional repertoire above the threshold focuses mainly on the COG J, with just a few interesting hubs belonging to COGs I, K and L.

Indeed, the PPI network of each bacterial species is characterized by a highly connected core of conserved ribosomal proteins, the components of multi-subunit complexes whose corresponding genes are mostly essential [31, 35] and code for supra-molecular complexes, that pile up in the bump we have observed for the degree distribution (figure1). Hence, what we are seeing here is essentially the ribosome, and related protein complexes such as RNA Polymerase. Indeed, the ribosome is the only molecular machine in bacteria in which a given protein could legitimately have 40 or more protein binding partners, with the help of rRNA mediating interactions [42].

We believe that the observations we have presented here can have implications both for the prediction of gene essentiality, based on the knowledge of PPI networks, and for the prediction of interactions between proteins, based on genetic information [43, 44]. It is interesting to note that our results are consistent with a previous study based on inferred bacterial co-conserved networks based on phylogenetic profiles [35]. The coupled flows of information from the genetic level up to the proteomic level and vice-versa should be further systematically investigated, taking into account archaeal and prokaryotic genomes in the search for emerging multi-layer structures that could offer basic theoretical grounds for clinical and systemic applications, for instance related to antimicrobial resistances [45–48].

